# A synthetic mechanogenetic gene circuit for autonomous drug delivery in engineered tissues

**DOI:** 10.1101/2020.04.29.069294

**Authors:** Robert J. Nims, Lara Pferdehirt, Noelani B. Ho, Alireza Savadipour, Jeremiah Lorentz, Sima Sohi, Jordan Kassab, Alison K. Ross, Christopher J. O’Conor, Wolfgang B. Liedtke, Bo Zhang, Amy L. McNulty, Farshid Guilak

## Abstract

Mechanobiologic signals regulate cellular responses under physiologic and pathologic conditions. Using synthetic biology and tissue engineering, we developed a mechanically-responsive bioartificial tissue that responds to mechanical loading to produce a pre-programmed therapeutic biologic drug. By deconstructing the signaling networks induced by activation of the mechanically-sensitive ion channel transient receptor potential vanilloid 4 (TRPV4), we created synthetic TRPV4-responsive genetic circuits in chondrocytes. We engineered these cells into living tissues that respond to mechanical loading by producing the anti-inflammatory biologic drug, interleukin-1 receptor antagonist. Chondrocyte TRPV4 is activated by osmotic loading and not direct cellular deformation, suggesting tissue loading is transduced into an osmotic signal that activates TRPV4. Either osmotic or mechanical loading of tissues transduced with TRPV4-responsive circuits protected constructs from inflammatory degradation by interleukin-1α. This synthetic mechanobiology approach was used to develop a mechanogenetic system to enable long-term, autonomously regulated drug delivery driven by physiologically-relevant loading.

## Introduction

Smart biomaterials or bioartificial tissues that autonomously respond to biologic cues and drive a therapeutic or restorative response are promising technologies for treating both chronic and acute diseases (*1*). Mechanotherapeutics in particular, are a rapidly growing class of smart biomaterials that use mechanical signals or mechanical changes within diseased tissues to elicit a therapeutic response (*2-4*) and ameliorate the defective cellular mechanical environment (*5-8*). Current mechanotherapeutic technologies rely on exogeneous protein drug delivery or ultrasound stimulation, or synthetic polymer implants that offer a finite lifespan for drug delivery (*9-11*). Creating systems with cellular-scale resolution of mechanical forces that offer long-term, feedback-controlled synthesis of biologic drugs could provide a completely new approach for therapeutic delivery.

In contrast to synthetic polymers, biological tissues grow, adapt, and respond to mechanobiologic signals through the use of specialized molecular components, such as mechanically-sensitive ion channels and receptors that transduce specific stimuli from the physical environment (*12, 13*). In particular, mechanosensitive ion channels are sensitive to both context and deformation mode, making them uniquely suited as mechanotherapeutic sensors (*14-16*). The transient receptor potential (TRP) family is a class of selective ion channels including some mechanically-sensitive members, such as TRPA1, TRPV1, and TRPV4 (*17, 18*). TRPV4 is activated by osmotic stress and plays an important role in the mechanosensitivity of various tissues such as articular cartilage, uterus, and skin (*19-23*). In cartilage, TRPV4 has been shown to regulate the anabolic biosynthesis of chondrocytes in response to physiologic mechanical strain (*24*).

Osteoarthritis is a chronic joint disease for which there are no available disease modifying drugs, ultimately leading to a total joint replacement once the diseased and degraded cartilage and surrounding joint tissues incapacitate a patient from pain and a loss of joint function (*25*). Cartilage tissue engineering is a promising strategy to resurface damaged and diseased articular cartilage with an engineered cartilage tissue construct as a means to reduce the need for, or prolong the time before, a total joint replacement (*26, 27*). An ongoing challenge in the field, however, is developing engineered cartilage constructs that withstand both the high mechanical loads present within the articular joint and the chronic inflammation present within an osteoarthritic joint (*28, 29*). For example, the knee cartilage of healthy individuals can experience compressive strains of ∼5-10% during moderate exercise, while individuals with a history of joint injury or high body mass index, populations at risk for developing osteoarthritis, can experience higher magnitudes of cartilage compression under similar loading (*30-32*). In this regard, a bioartificial tissue that can synthesize biologic drugs in response to inflammation or mechanical loading, either independently or concurrently, could greatly enhance the therapeutic potential of an engineered tissue replacement.

In this study, we engineered a mechanically-responsive bioartificial tissue construct for therapeutic drug delivery by using the signaling pathways downstream of the mechano/osmosensitive ion channel TRPV4 to drive synthetic mechanogenetic gene circuits (Fig. 1). To do this, we first establish that mechanical loading of tissue engineered cartilage activates TRPV4 through fluctuations in the local osmotic environment and not direct mechanical deformation of chondrocytes. We next deconstructed the gene regulatory networks and signaling pathways evoked by TRPV4 activation and revealed the transient activation of several mechanosensitive transcription factors. We engineered synthetic gene circuits to respond to mechanical TRPV4-activation for driving transgene production of an anti-inflammatory molecule, interleukin-1 receptor antagonist (IL-1Ra). While IL-1Ra (drug name anakinra) is approved as a therapy for rheumatoid arthritis and has successfully attenuated osteoarthritis progression in pre-clinical models, clinical trials of IL-1Ra therapy patients with established osteoarthritis have not shown efficacy, suggesting controlled long-term delivery may be necessary for disease modification (*33-35*). Here, we show that mechanical or osmotic loading of implantable engineered cartilage tissue constructs induces an autonomous mechanogenetic response and protects tissues constructs from inflammatory insult, suggesting a modality for long-term *in vivo* drug delivery.

**Fig. 1.**
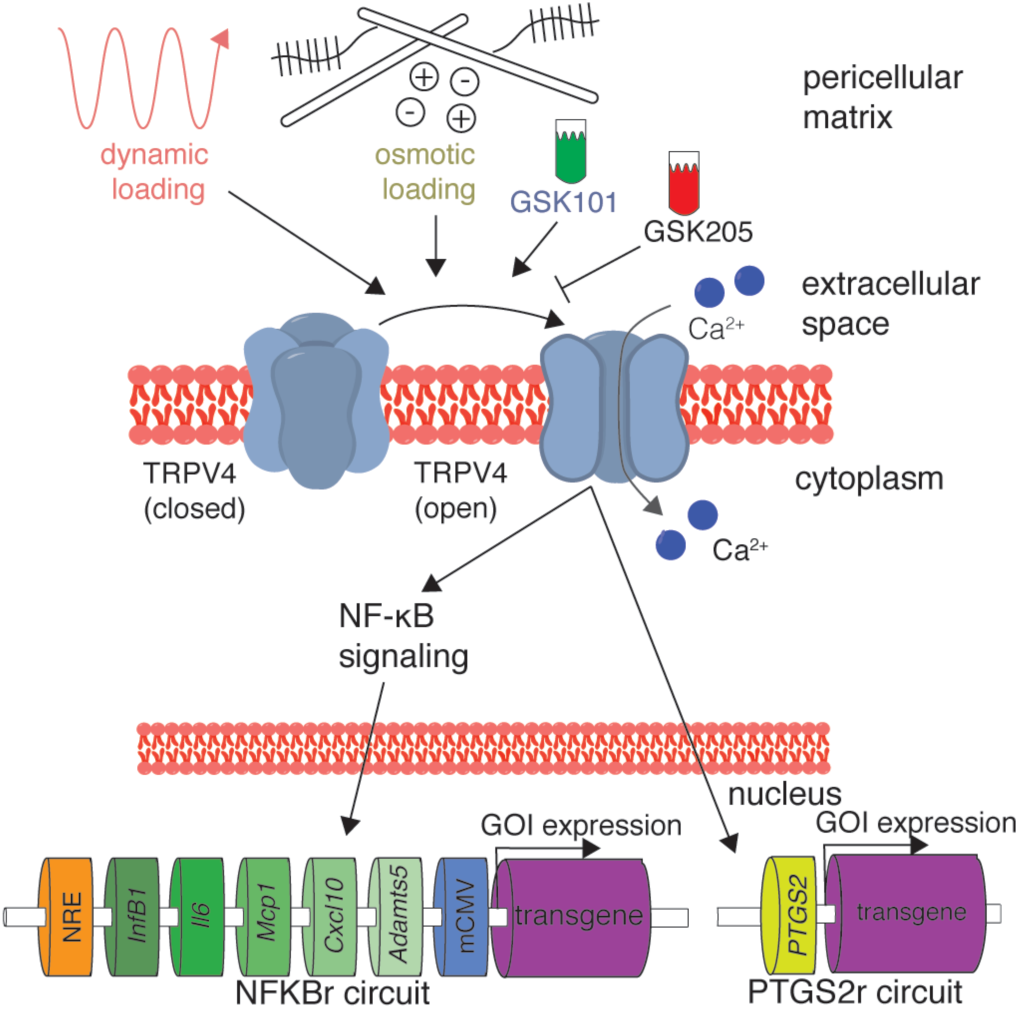
Mechanogenetic transduction and therapeutic drug delivery approach. Transient receptor potential vanilloid 4 (TRPV4) is an osmotically-sensitive cation channel in the cell membrane of chondrocytes, which can be activated by mechanical loading secondary to mechano-osmotic coupling through the extracellular matrix or pharmacologically with the agonist GSK101. TRPV4 can also be inhibited with the antagonist GSK205. Upon TRPV4 activation, chondrocytes respond with intracellular calcium signaling that initiates NF-*κ*B signaling and upregulation of the *PTGS2* gene. By lentivirally transducing synthetic mechanogenetic circuits that respond to either NF-*κ*B activation or *PTGS2* upregulation into chondrocytes within an engineered cartilage tissue, mechanically activated TRPV4 signaling was used to drive transgene production of either a luciferase reporter or the therapeutic anti-inflammatory biologic interleukin-1 receptor antagonist (IL-1Ra).

## Results

### The mechano-osmotic response of chondrocytes to loading is regulated by TRPV4

To deconstruct the mechanotransduction pathways through which chondrocytes perceive mechanical loading, we encapsulated freshly isolated primary porcine chondrocytes within an agarose hydrogel scaffold to engineer cartilage tissue constructs. Constructs were cultured for three weeks to allow extracellular matrix deposition before applying compressive loading and simultaneously imaging the intracellular calcium levels of the chondrocytes. This engineered cartilage system allows for mechanical signals to be transduced to chondrocytes through *de novo* synthesized extracellular matrix in a manner similar to *in vivo* mechanotransduction (*36-38*). Chondrocytes express an array of mechanically-sensitive ion channels and receptors, including TRPV4, PIEZO1 and PIEZO2, integrins, and primary cilia (*39*), so we investigated the mechanism by which physiologic magnitudes of dynamic mechanical loading are transduced by chondrocytes. Dynamic mechanical loading of engineered cartilage constructs immediately provoked a 108% increase of intracellular calcium response compared with unloaded tissues (p=0.022, Fig. 2a). Inhibiting TRPV4 activity using the TRPV4 antagonist GSK205 suppressed calcium signaling (p=0.03, GSK205 supplementation reduced signaling ∼47%), implicating TRPV4 as a critical component in transducing compressive loading in chondrocytes.

**Fig. 2.**
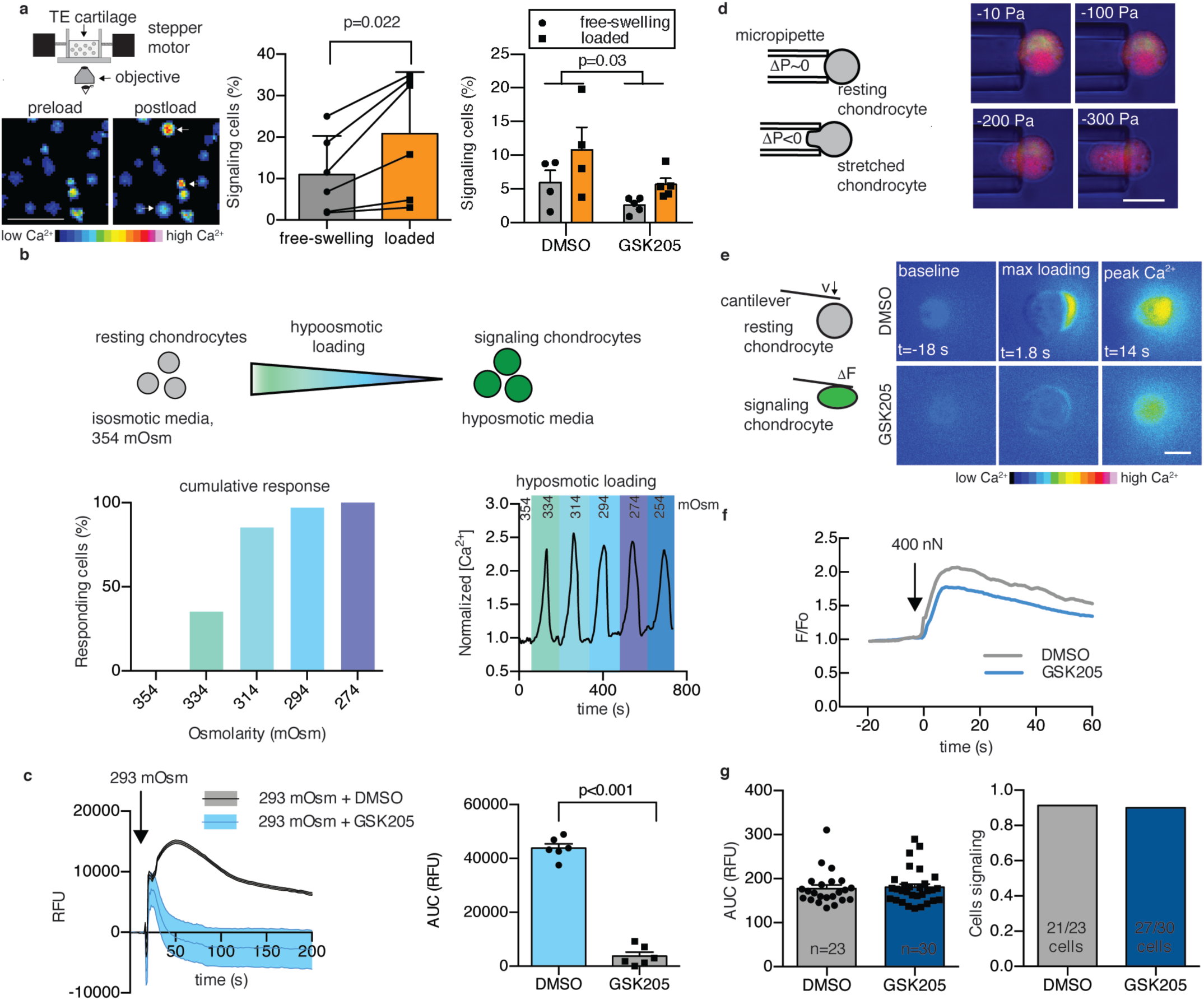
Mechanical responsiveness of chondrocytes is mediated by hypo-osmotic stimulation of TRPV4. (a) Set-up of real-time cellular imaging of mechanical loading. Loading chondrocytes within engineered cartilage increases intracellular calcium compared to free swelling. Arrows indicate immediately-responsive cells. Scale bar is 50 µm. The number of cells exhibiting intracellular calcium signaling increased by 108% after loading, and GSK205 suppressed cellular calcium signaling (n=4-5 constructs per treatment). (b) Isolated chondrocytes are sensitive to osmotic perturbations and exhibit intracellular calcium increases in response to hypo-osmotic stimulation (n=15-20 cells per group, calcium response is normalized to calcium levels at 354 mOsm). (c) Chondrocyte responsiveness to hypo-osmotic stimulation is inhibited with GSK205 (n=6 per treatment). (d) Chondrocytes are not sensitive to direct membrane stretch applied under iso-osmotic, iso-volumetric conditions with micropipette aspiration (n=38, scale bar is 10 µm). (e) Direct cellular compression under a 400 nN load with an atomic force microscope induces intracellular calcium signaling. Scale bar is 10 µm. (f) GSK205 does not modulate calcium response of chondrocytes to AFM loading. (g) TRPV4 inhibition alters neither the intensity of calcium responsiveness nor the population response to AFM compression (n=23-30 cells per group). Data are presented as mean ± SEM.

While the molecular structure of TRPV4 has recently been reported (*40*), the mechanisms underlying TRPV4 activation are complex and may involve direct mechanical activation or osmotic activation secondary to mechanical loading of the charged and hydrated extracellular matrix (*17, 21, 41, 42*). To determine the physical mechanisms responsible for TRPV4 activation secondary to tissue compression, we subjected freshly isolated chondrocytes to osmotic loading, direct membrane stretch, and direct single-cell mechanical compression (Fig. 2b, d, e). To better understand the biophysical mechanisms underlying TRPV4 activation, we determined the mechanical state of the chondrocyte membrane in each of these cases using finite element modeling (FEBio; www.febio.org) (*43*). Chondrocytes rapidly responded to physiologically-relevant changes in media osmolarity through intracellular calcium signals (Fig. 2b). Calcium signaling was highly sensitive to modest changes in osmolarity, with an osmolarity shift of −20 mOsm inducing intracellular signaling in 35% of chondrocytes and −80 mOsm inducing activation of 100% of chondrocytes. Inhibition of TRPV4 with GSK205 suppressed calcium signaling caused by hypo-osmotic loading (Fig. 2c) or by the TRPV4 agonist GSK1016790A (GSK101) (Supplemental Fig. S1). Chondrocyte volumetric analysis and finite element modeling of hypo-osmotic stress showed that a change of −60 mOsm, sufficient to induce signaling in nearly all chondrocytes, increased cellular volume by 13% and induced an apparent first principle strain of 0.04 homogeneously throughout the membrane (Supplemental Figs. S2 and S3). To investigate the role of direct membrane stretch in TRPV4 activation in the absence of osmotic fluctuations, we used micropipette aspiration of individual chondrocytes to apply controlled deformation of the cell membrane (Fig. 2d). Surprisingly, micropipette aspiration did not provoke any calcium signaling response in chondrocytes, with only 1 of 38 tested cells responding with increased intracellular calcium, indicating that membrane deformation *per se* was not the primary signal responsible for mechanical activation. Finite element simulations of the micropipette experiment showed the presence of heterogenous apparent membrane strains in aspirated chondrocytes, with a first principle strain reaching 0.31 around the micropipette mouth and ∼0.04 within the micropipette under an applied pressure of 100 Pa (Supplemental Fig. S3e-h). To test whether calcium signaling in response to direct mechanical compression of chondrocytes is mediated by TRPV4, we loaded isolated chondrocytes with an atomic force microscope (AFM) to 400 nN (*44*). Only at high, pathologic levels of mechanical compression did direct chondrocyte compression provoke intracellular calcium signaling, but this response was not inhibited by GSK205 (Fig. 2e-g). Finite element modeling predicted high membrane strains with an ultimate apparent first principle membrane strain of 2.27 around the cell at the peak cell deformations necessary to elicit intracellular calcium signaling (Supplemental Fig. S3i-l). Together, these findings suggest that activation of TRPV4 in chondrocytes *in situ* may not result directly from cellular strain but rather from osmotic fluctuations induced from the deformation of a chondrocyte’s osmotically active environment. Due to the complex relationship between deformational compressive loading of engineered cartilage and osmotic loading, we will hereafter refer to deformational compressive loading (concomitant with the secondary osmotic effects) as mechanical loading and direct changes in the media osmolarity as osmotic loading.

### TRPV4 activation in chondrocytes induces transient anabolic and inflammatory signaling networks

We next sought to understand the time course of specific downstream signaling pathways and gene regulatory networks regulated by mechanical activation of TRPV4 using microarray analysis. As TRPV4 is a multimodal channel, downstream signaling is likely dependent on the activation mode, as well as cell and tissue type. For these studies, primary porcine chondrocytes were cast in agarose to create engineered cartilage constructs, and subjected to either compressive mechanical loading (10% sinusoidal peak-to-peak strain at 1 Hz for 3 h as described in (*45*)), or pharmacologic stimulation with the TRPV4 agonist GSK101 (1 nM for 3 h), or left unloaded (free-swelling controls), and measured the transcriptional activity at 0, 3, 12, and 20 h following an initial round of stimulation and then at 24 and 72 h after additional daily bouts of mechanical loading or GSK101 (n=3 per condition per time point, Fig. 3a). In the first 20 h following initial stimulation there were 43 transcripts upregulated under both mechanical loading and pharmacologic stimulation (Fig. 3b and Supplementary Tables 1 and 2) compared to unloaded controls. Upregulation of these targets was immediate and subsided quickly, with transcript levels returning to baseline 12 to 20 h after activation. In particular, the cAMP/Ca^2+^-responsive transcription factors *C-JUN, FOS, NR4A2*, and *EGR2* (Fig. 3c) were all upregulated in response to TRPV4-mediated calcium signaling.

**Fig. 3.**
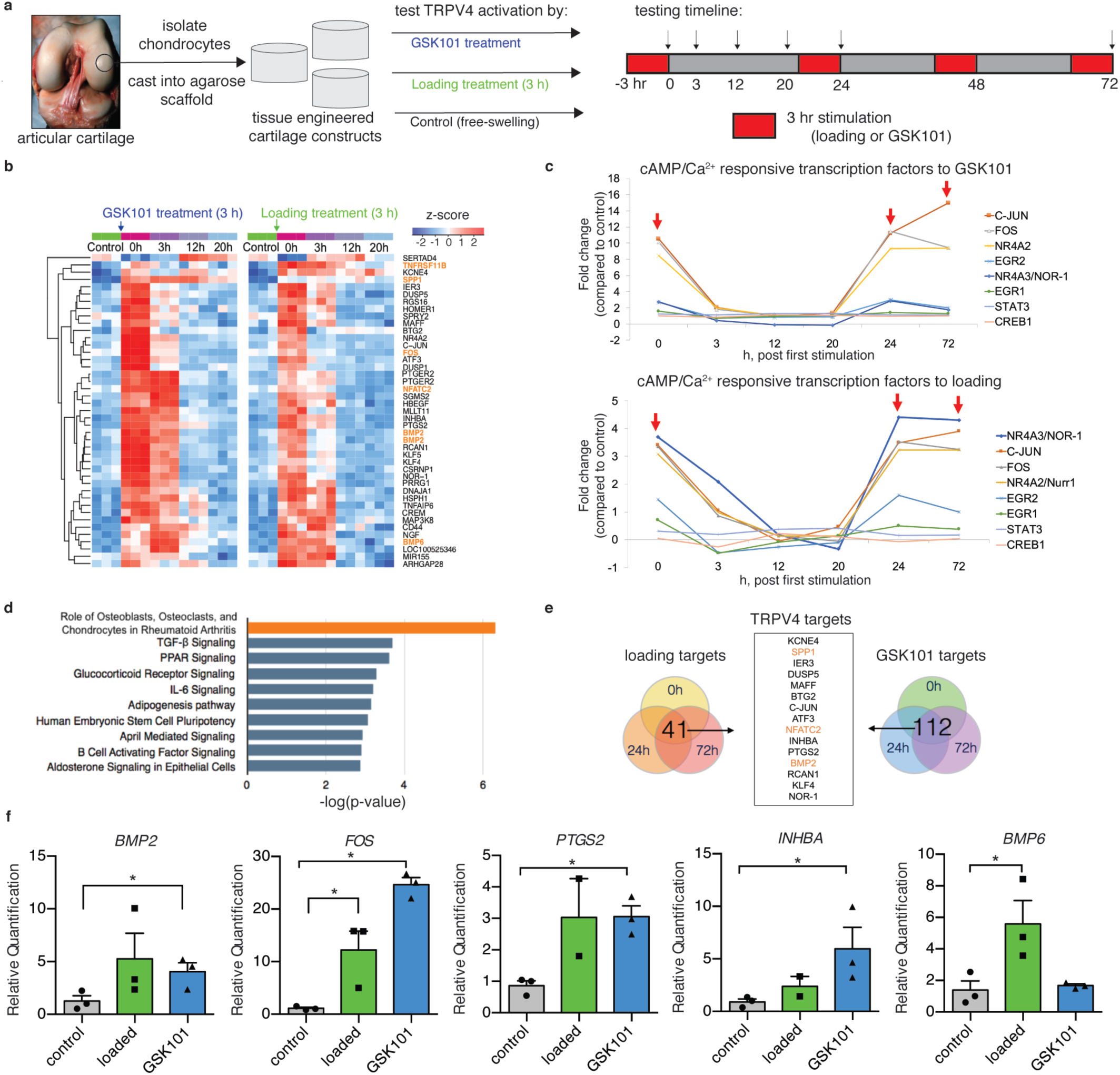
Transcriptomic profile induced by TRPV4 activation. (a) Engineered cartilage tissue constructs were made from isolated primary porcine chondrocytes cast into agarose hydrogels. Tissue constructs were cultured in nutrient-rich media before deformational mechanical loading or GSK101 pharmacologic stimulation (red, 3 h/round) following the indicated time course and cartilage construct harvest (arrows). (b) 41 genes were differentially upregulated in response to TRPV4 activation and levels returned back to baseline after 12-20 h post loading (n=3/treatment/timepoint). (c) cAMP and calcium-responsive transcription factors were immediately and highly regulated by both mechanical loading and GSK101 stimulation (red arrows indicate removal from loading). (d) Pathway analysis based on transcription activity suggests both inflammatory and anabolic pathways are strongly regulated by TRPV4 activation. (e) Analysis of gene targets response after all bouts of mechanical loading and all bouts of GSK101 stimulation produces a list of distinctly-TRPV4 sensitive genes. (f) TRPV4-responsive targets from the microarray analysis were confirmed by qPCR (n=2-3) a one-tailed t-test was used to test whether loaded or GSK101 groups were significantly upregulated with treatment.

We then used Ingenuity Pathway Analysis to identify candidate targets for a synthetic gene circuit that would be responsive to TRPV4 activation. The most significantly enhanced pathway was “the role of osteoblasts, osteoclasts, and chondrocytes in rheumatoid arthritis,” with members including transcription factors (*FOS* and *NFATC2*), extracellular matrix synthesis gene (*SPP1*), growth factors (*BMP2* and *BMP6*), and *TNFSF11*, the decoy ligand for RANKL, all of which were upregulated by TRPV4 activation (Fig. 3d). The inflammatory pathways “IL-6 signaling” and “B cell activating factor signaling” were also significantly upregulated by TRPV4 activation, as were the established chondrocyte anabolic pathways “TGF-β signaling” and “Glucocorticoid signaling.” Due to the rapid response times of nearly all of the differentially expressed genes, we further analyzed which genes responded to repeated bouts of TRPV4 stimulation at all three time points (0, 24, and 72 h). There were 41 transcripts repeatedly responsive to mechanical loading and 112 transcripts repeatedly responsive to GSK101 supplementation. Of these targets, 15 genes were commonly responsive to both mechanical loading and pharmacologic activation of TRPV4 (Fig. 3e) and include both pathways associated with inflammation (*PTGS2, SPP1, ATF3*) and cartilage development and homeostasis (*BMP2, DUSP5, NFATC2, INHBA*). The most robustly regulated and cartilage-relevant genes were confirmed by qPCR (Fig. 3f). Taken together, these data suggest that the TRPV4 mechanotransduction pathway involves activation of a rapidly resolving inflammatory pathway as part of the broad anabolic response to physiologic mechanical loading.

### Synthetic mechanogenetic circuits respond to TRPV4 activation to drive transgene expression

TRPV4 activation in response to mechanical loading upregulates a diverse group of targeted genes. Based on the TRPV4-activated signaling pathway (Supplemental Fig. S4), we identified activation of the nuclear factor kappa-light chain enhancer of activated B cells (NF-*κ*B) pathway and upregulation of the prostaglandin-endoperoxide synthase 2 (*PTGS2*) gene as two distinct avenues to construct TRPV4-responsive synthetic mechanogenetic gene circuits. Targeting the NF-*κ*B signaling pathway and regulation of the *PTGS2* promoter (Fig. 1), we developed two lentiviral systems that would either (1) respond to NF-*κ*B activity by linking 5 synthetic NF-*κ*B binding motifs and a NF-*κ*B negative regulatory element (*46*) with the cytomegalovirus (CMV) enhancer to drive transgene expression of either the therapeutic anti-inflammatory biologic IL-1Ra or a luciferase reporter (henceforth referred to as NFKBr-IL1Ra and NFKBr-Luc, respectively) or (2) respond to *PTGS2* regulation by using a synthetic human *PTGS2* promoter to drive either IL-1Ra or luciferase expression (henceforth referred to as PTGS2r-IL1Ra and PTGS2r-Luc, respectively). We then created mechanogenetically-sensitive engineered cartilage tissue constructs by lentivirally-transducing primary porcine chondrocytes with a mechanogenetic circuit and seeding these cells into an agarose hydrogel to produce synthetically-programmed cartilage constructs (Fig. 4a).

**Fig. 4.**
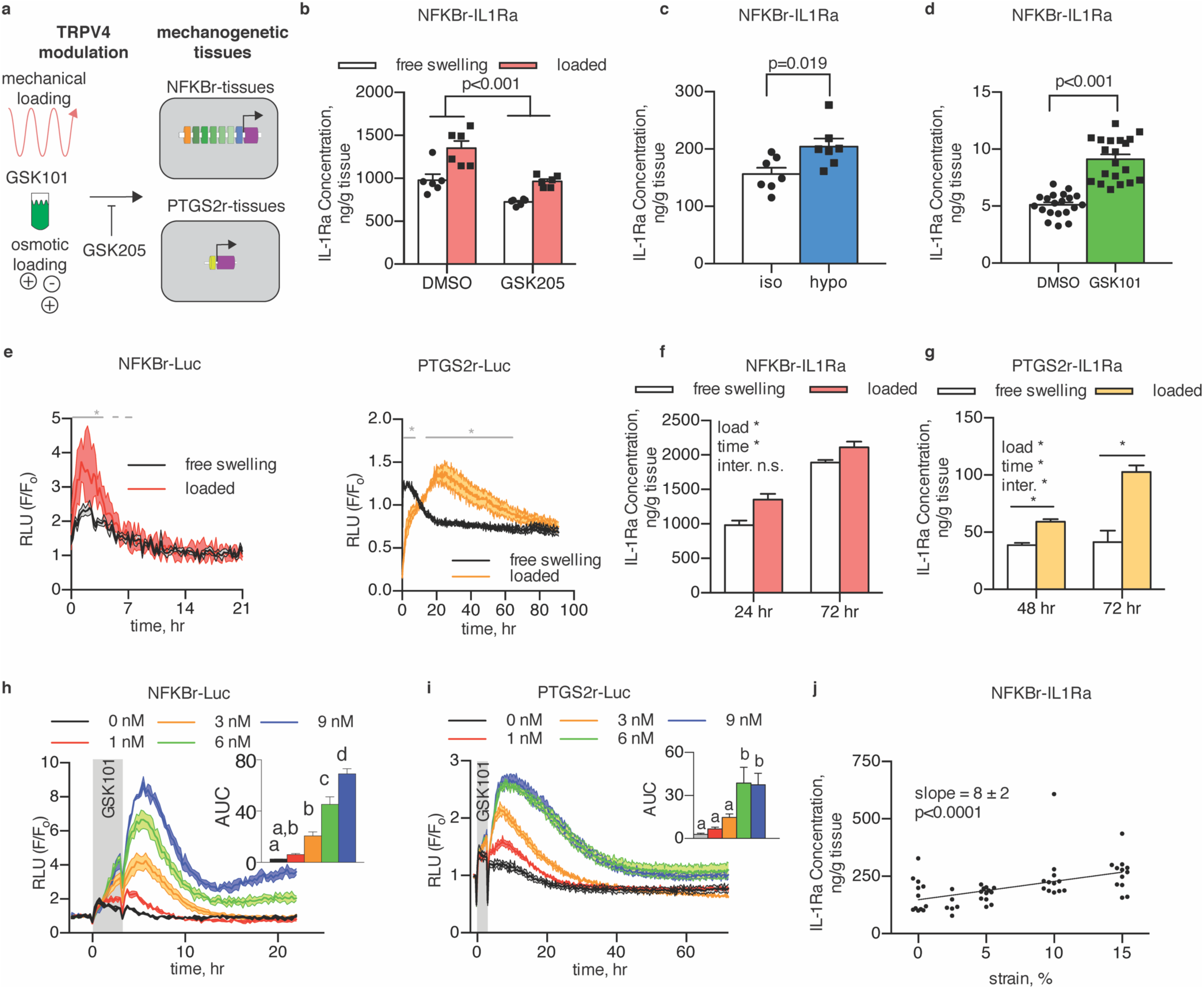
Mechanogenetic constructs respond to TRPV4 activation. (a) Mechanical loading, osmotic loading, or GSK101 stimulation was applied to mechanogenetic tissues; GSK205 inhibit TRPV4 activation. (b) NFKBr-IL1Ra tissues respond to mechanical loading through increased IL-1Ra (p<0.001). IL-1Ra is reduced with GSK205 supplementation (p<0.001, n=6 per treatment). (c) Exposure of NFKBr-IL1Ra to hypo-osmotic media produces more IL-1Ra than iso-osmotic media exposure (p=0.019, n=7 per group). (d) NFKBr-IL1Ra tissues exposed to GSK101 stimulation produce more IL-1Ra than vehicle controls (p<0.001, n=20 per group). (e) Mechanical loading of NFKBr-Luc tissues quickly activates and inactivates circuits, while PTGS2r-Luc tissues take longer to reach the peak and return to baseline (gray line denotes p<0.05 between free-swelling and load, n=3-6 per group). (f) NFKBr-IL1Ra tissue response to loading after 24 h and 72 h, indicating differential expression in first 24 h. (g) PTGS2r-IL1Ra tissues respond to loading through 72 h. (h) NFKBr-Luc tissues respond dose-dependently to TRPV4 activation via GSK101 through 9 nM GSK101 (p<0.05, n=2-4 per group). (i) PTGS2r-Luc tissues are sensitive to GSK101 up to 6 nM (p<0.05, n=2-4 per group). (j) NFKBr-IL1Ra tissue response is dose-dependent to compressive mechanical loading strain from 0 to 15 % (p<0.001, n=5-12). Data are presented as mean ± SEM.

To test whether mechanogenetic tissues actively respond to mechanical stimulation for transgene production, we first applied compressive mechanical loading to NFKBr-IL1Ra-transduced mechanogenetic cartilage constructs, which showed a 38% increase in IL-1Ra production (p=0.006) with mechanical loading (Fig. 4b) compared to free swelling controls. To establish whether this response was dependent on TRPV4, we antagonized TRPV4 with GSK205 and observed a significant attenuation of NFKBr-IL1Ra circuit activity (Fig. 4b), demonstrating the mechanogenetic circuit responded to both mechanical loading (p<0.001) and TRPV4 antagonism (p<0.001). Interestingly, GSK205 also reduced circuit activation in unloaded tissues, suggesting TRPV4 activation in chondrocytes may not be entirely dependent on mechanical loading. To further assess the specificity of TRPV4-regulation in mechanogenetic cartilage constructs, we applied direct hypo-osmotic loading and pharmacologic GSK101 stimulation to NFKBr-IL1Ra tissues. A step change to hypo-osmotic media (−200 mOsm change) activated the NFKBr-IL1Ra mechanogenetic circuit (p=0.019, Fig. 4c) compared with an iso-osmotic media change (0 mOsm change). Additionally, daily pharmacologic activation of TRPV4 with 1 nM GSK101 for 3 h/d over the course of 5 days activated the NFKBr-IL1Ra circuit compared to DMSO controls (p<0.001, Fig. 4d). Engineered cartilage constructs responded repeatedly and reproducibly to the daily GSK101 stimulation (Supplemental Fig. S5), demonstrating the role of TRPV4 in activating the mechanogenetic NFKBr-circuits and confirming that a cell-based mechanotherapy can offer prolonged and unabating biologic drug delivery. Testing of conditioned media from constructs seeded with either non-transduced chondrocytes or chondrocytes transduced with a green fluorescent protein (GFP)-expression cassette found that only cells transduced with a mechanogenetic IL-1Ra-producing circuits were capable of synthesizing detectable levels of IL-1Ra.

To determine the sensitivity and temporal response kinetics of our engineered tissue system, factors critical for the effectiveness of any drug delivery system, we used bioluminescence imaging of NFKBr-Luc or PTGS2r-Luc constructs to determine the dynamic response of mechanogenetic cartilage constructs to TRPV4 activation by mechanical loading or GSK101 pharmacologic stimulation. In response to 10% compressive mechanical loading (Fig. 4e), NFKBr-Luc tissue constructs rapidly peaked (1.8 ± 0.2 h to peak) and decayed (T_50%_ decay time = 3.4 ± 0.4 h) with loaded samples returning to baseline 4 h after loading. PTGS2r-Luc tissue constructs were slower to activate (21.7 ± 2.7 h to peak) and remained activated for a longer duration (T_50%_ decay time = 22.3 ± 1.7 h). This differential in time delivery kinetics may provide strategies by which mechanical loading inputs can drive both short- and long-term drug production by judicious mechanogenetic circuit selection in a single therapeutic tissue construct. To test whether IL-1Ra production followed similar differential production rates, we mechanically loaded NFKBr-IL1Ra and PTGS2r-IL1Ra tissues constructs and measured protein levels of IL-1Ra released in the media. One round of mechanical loading activated NFKBr-IL1Ra tissues by 24 h, as measured by an increase in IL-1Ra produced by loaded tissue constructs compared to unloaded constructs (an increase in IL-1Ra production of 372 ± 265 ng/g, mean ± 1 s.d.). This differential in IL-1Ra concentration between loaded and free swelling constructs remained unchanged by 72 h (loaded tissues produced 220 ± 223 ng/g more than unloaded constructs at this time point), indicating loaded NFKBr-tissues were not continually activated and resumed baseline activity levels after 24 hours (Fig. 4f). Conversely, in a preliminary experiment, a single round of mechanical loading did not differentially activate PTGS2r-IL1Ra in the 24 h after loading (Supplementary Fig. S6). Informed by the bioluminescent imaging, we measured conditioned media of PTGS2r-IL1Ra tissues 48 and 72 h after a single round of mechanical loading, however, and found tissues produced more IL-1Ra after 48 h compared to free swelling tissues (difference in means of 21 ng/g), and the IL-1Ra difference between loaded and free swelling tissues continued to increase by 72 h (difference in means of 61 ng/g). This increased effect size confirmed mechanically loaded PTGS2r-tissue constructs remained activated up to 72 h after loading and demonstrated a longer-acting mechanogenetic response in the PTGS2r-system (Fig. 4g). Based on the bioluminescent imaging results, we did not measure IL-1Ra levels past 72 h. These results show that complex drug delivery strategies can be programmed into a single mechanogenetic tissue constructs to produce multiple modes and timescales of therapeutic or regenerative biologic drug delivery.

For insight into the dose-response relationship and sensitivity of the mechanogenetic gene circuits, we imaged NFKBr-Luc and PTGS2r-Luc bioluminescence in response to different doses of GSK101. Temporal imaging revealed that GSK101 stimulation produced similar rise and decay kinetics as TRPV4-activation from mechanical loading (Fig. 4e) in NFKBr-Luc (Fig. 4h) and PTGS2r-Luc (Fig. 4i) cartilage tissue constructs compared to unstimulated controls. Both mechanogenetic tissue constructs displayed a clear dose-dependent activation from GSK101 stimulation as demonstrated by the AUC. NFKBr-Luc tissue constructs were responsive from 1 through 9 nM GSK101, whereas PTGSr-Luc tissue constructs plateaued in responsiveness at 6 nM GSK101. Based on this pharmacologic sensitivity, we hypothesized mechanogenetic constructs would be increasingly activated by higher mechanical loading strains through increased osmotic stimulation (Fig. 2b). As NFKBr-Luc tissues demonstrated the most pronounced GSK101 dose-dependent response, we applied dynamic compressive strain amplitudes from 0 to 15% to NFKBr-IL1Ra tissue constructs to span the range of physiologic strains expected for articular cartilage *in vivo* (*36*). We measured elevated production of IL-1Ra with increasing compressive strains (Fig. 4j; slope = 8 ± 2 ng/g IL-1Ra per %strain, p<0.001), demonstrating that drug production is directly responsive to the magnitude of mechanical loading in our mechanogenetic engineered tissues.

### Mechanogenetic engineered cartilage activation protects cartilage tissues from IL-1α-induced inflammation-driven degradation

The long-term success of engineered cartilage implants depends on the ability of implants to withstand the extreme loading and inflammatory stresses within an injured or osteoarthritic joint (*47*). We hypothesized that TRPV4 activation would activate mechanogenetic tissue constructs to produce therapeutic levels of IL-1Ra and protect engineered cartilage constructs and the surrounding joint from destructive inflammatory cytokines. As our mechanogenetic circuits rely on signaling pathways that overlap with the cellular inflammatory response, we examined the dose-response of IL-1Ra production in response to the inflammatory cytokine interleukin-1α (IL-1α; Figs. 5a and c). NFKBr-IL1Ra mechanogenetic constructs responded to exogeneous IL-1α supplementation following a dose-dependent characteristic, consistent with findings of IL-1-induced NF-*κ*B signaling in chondrocytes ((*48*), Fig. 5b). Importantly, this dose-dependent response to IL-1α was present up to 10 ng/mL and offered prolonged and robust production of IL-1Ra, promoting the notion that cell-based tissues may offer a more sustained ability to produce therapeutic biomolecules relative to traditional acellular approaches (Supplementary Fig. S7). Mechanical loading further enhanced IL-1Ra production in the presence of IL-1α, demonstrating that even in the presence of high levels of inflammation, mechanical loading further potentiates NF-*κ*B signaling (Supplementary Fig. S8). Interestingly mechanogenetic PTGS2r-IL1Ra and PTGS2r-Luc constructs demonstrated no sensitivity to IL-1α (Figs. 5d and e). Importantly, and in contrast to NFKBr-tissue constructs, the selective sensitivity to TRPV4 activation and not to IL-1α in PTGS2r-tissue constructs suggests that this system is distinctively sensitive to mechano- or osmo-activation of TRPV4. This distinction was further demonstrated in tissues exposed to both IL-1α inflammation and osmotic loading, wherein only loading played a significant influence (Supplemental Fig. S9, p=0.0002 for loading and p=0.14 for inflammation).

**Fig. 5.**
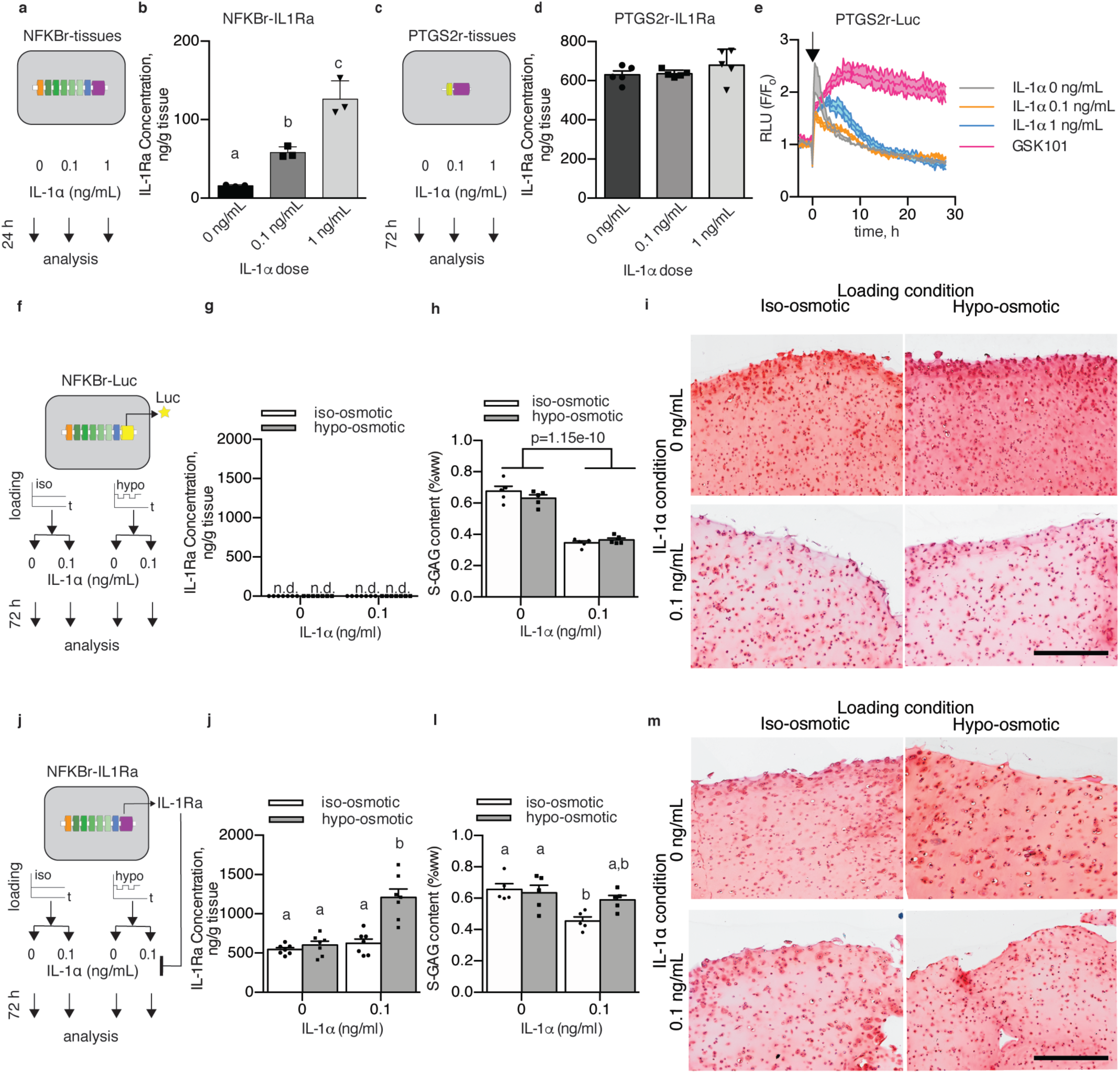
Osmotic loading of mechanogenetic constructs protects against IL-1α. (a) The inflammatory response of NFKBr-IL1Ra constructs under IL-1α supplementation. (b) NFKBr-IL1Ra tissues produce IL-1Ra in response to IL-1α (n=3, p<0.001). (c) The inflammatory response of PTGS2r-IL1Ra tissues under IL-1α supplementation. (d) PTGS2r-IL1Ra tissues do not respond to IL-1α (n=5). (e) PTGS2r-Luc tissues are not altered by IL-1α by are modulated by chronic GSK101 (n=2 per condition, arrow indicates stimulation). (f) Osmo-inflammatory response of NFKBr-Luc tissues using osmotic loading (3 h/d) and IL-1α (0 or 0.1 ng/mL) applied to NFKBr-Luc tissues. (g) NFKBr-Luc tissues do not produce IL-1Ra (n=7). (h) NFKBr-Luc tissues lost S-GAG in the presence of IL-1α (n=5 per group). (i) Histological reduction of safranin-O staining with IL-1α supplementation. (j) Osmo-inflammatory response of NFKBr-IL1Ra tissues using osmotic loading (3 h/d) and IL-1α (0 or 0.1 ng/mL). (j) IL-1Ra was increased with inflammation and osmotic loading (n=7, p<0.001). (l) S-GAG content in NFKBr-IL1Ra tissues with IL-1α supplementation (0 or 0.1 ng/mL) and/or osmotic loading (different letters denote significant differences, p<0.05, n=5). (m) NFKBr-IL1Ra tissues displayed similar safranin-O staining without IL-1α supplementation, while IL-1α supplementation reduced safranin-O staining in iso-osmotic tissues but not hypo-osmotic tissues. Data are presented as mean ± SEM.

To test whether TRPV4 activation-induced IL-1Ra production would protect engineered mechanogenetic tissue constructs against IL-1α, we co-cultured mature, 21-day-old NFKBr-tissue constructs with articular cartilage explants in the presence of IL-1α inflammation and daily hypo-osmotic loading (incubation in 180 mOsm media for 3 h/d; standard iso-osmotic media was maintained 380 mOsm) to model the inflammation and osmotic conditions present in the cartilage of arthritic joints exposed to daily loading (Figs. 5f and j). Using NFKBr-Luc tissue constructs, which lack an anti-inflammatory response to TRPV4- or IL-1α-activation (Fig. 5g), to simulate how conventional engineered cartilage constructs would respond in an arthritic joint, constructs lost 45.6% of their sulfated-glycosaminoglycan content after 72 h of IL-1α treatment (Fig. 5h, S-GAG). S-GAGs are essential structural molecules that impart mechanical integrity and strength in both engineered and native cartilage (*49, 50*). Osmotic loading did not modulate this response and the substantial S-GAG loss was observable histologically through diminished safranin-O staining throughout the tissue (Fig 5i). In the anti-inflammatory NFKBr-IL1Ra tissues constructs, osmotic loading in the presence of inflammatory IL-1α increased IL-1Ra production 93% over iso-osmotic control tissues also cultured with IL-1α (Fig. 5j). After 72 h of treatment, IL-1α induced a 30.8% loss of S-GAG in NFKBr-IL1Ra engineered cartilage constructs under iso-osmotic conditions, while NFKBr-IL1Ra engineered cartilage that was incubated with IL-1α and osmotically loaded did not significantly lose their S-GAG content (Fig. 5l). Histologically, hypo-osmotic loading NFKBr-IL1Ra engineered cartilage constructs maintained rich safranin-O staining of S-GAG throughout the constructs, even in the presence of IL-1α (Fig. 5m). Interestingly, explant proteoglycan levels were similar across all IL-1α treatment groups. Together these data demonstrate engineered mechanogenetic cartilage constructs can produce anti-inflammatory IL-1Ra in response to osmotic loading at levels sufficient for engineered tissue protection in a proinflammatory environment mimicking the conditions present in an osteoarthritic joint (*51*).

## Discussion

By combining synthetic biology and tissue engineering, we developed a novel class of bioartificial material that is mechanogenetically-sensitive. This system functions by redirecting endogenous mechanically-sensitive ion channels to drive synthetic genetic circuits for converting mechanical inputs into programmed expression of a therapeutic transgene. By engineering these cells into a functional tissue construct, this system provides the potential to repair or resurface damaged cartilage, while providing site-specific, mechanically-induced anti-cytokine therapy against inflammation. Our approach is based on redirection of the downstream response to activation of the TRPV4 ion channel – a critical mechanosensor in cartilage – to transduce tissue-scale deformational mechanical loading via mechano-osmotic coupling in the charged extracellular matrix. By deconstructing the gene regulatory networks activated by mechanically-induced TRPV4, we co-opted chondrocyte-endogenous signaling machinery to drive synthetic circuits for the production of therapeutic biologic drugs, using the anti-inflammatory drug IL-1Ra as a model system for proof of concept. Our results also demonstrate the use of distinct signaling networks for defining the specificity, timing, and dose-response for the expression of therapeutic biologic drugs. We show that a single round of mechanical loading can induce short-term and long-term responses based on the particular response of the synthetic circuit. While we targeted a treatment for cartilage repair and osteoarthritis, pathologic mechanical loading and mechanosignaling play a role in a broad range of acute and chronic diseases (*12*), suggesting a wide range of potential therapeutic applications for mechanogenetically regulated cells or tissues requiring autonomous cellular control systems.

While we developed mechanogenetic cartilage tissues based on TRPV4-activation here, the use of other native, mechanically-sensitive ion channels and receptors provide an attractive source of mechanosensors that can be elicited to provoke synthetic outputs. Expanding mechanogenetic approaches to additional mechanosensors with applications to other tissues would increase the range of physical stimuli that synthetic circuits can respond to, but requires an in-depth understanding of both the mechanical contexts necessary for mechanosensor activation and the resulting downstream signaling pathways. The fact that primary chondrocytes possess an array of different mechanically-sensitive ion channels and receptors highlights the potential opportunities to layer mechanosensor-specific circuits and produce systems responsive to different mechanical inputs that drive specific synthetic outputs. Our analysis here demonstrates that physiologic (∼10%, 1 Hz) mechanical loading of engineered cartilage is converted to a mechano-osmotic signal that activates TRPV4, evidenced by the GSK205-inhibition of chondrocyte calcium signaling in response to mechanical loading or osmotic loading of engineered cartilage, but not to direct cellular deformation or membrane deformation. To this end, we examined the mechanoresponsiveness of TRPV4 signaling and developed two genetic circuits that respond to TRPV4 activation. It is of interest to note that while endogenous *PTGS2* regulation subsided within 24 h after mechanical loading, our PTGS2r-circuit remained activated up to 72 h after loading, highlighting the role endogenous mechanisms of gene regulation may play, which are likely absent in our synthetic circuits. In this regard, the use of alternate chondrocyte mechanosensors remains an attractive area of research. For instance, the lack of a TRPV4-dependent response to high-strain compression via AFM loading is consistent with our earlier finding that high, potentially injurious, cellular strains are transduced via the PIEZO family of ion channels (*44*). The multimodal nature of TRPV4, and the TRP family more generally (*52*), suggests additional genetic circuits may be developed for alternate activation modes by characterizing the downstream signaling networks that respond to temperature or biochemical activation of the channel (*53*). Moreover, engineering novel mechanosensors may open the opportunity for developing custom mechanical activation modes and orthogonal downstream signaling in future mechanogenetic systems.

Our findings offer a detailed perspective of the complex cellular events initiated by TRPV4 activation in chondrocytes. TRPV4 has been thought to play a largely anabolic role in chondrocytes through enhanced synthesis of matrix molecules, S-GAG and collagen, and upregulation of TGF-β3 (*24*), which is evident in our GO/KEGG pathways hits of “TGF-β Signaling” and “Glucocorticoid Receptor Signaling”. However, our findings also revealed the presence of an acute, transiently resolving inflammatory response (pathways including “Chondrocytes in RA,” “IL-6 Signaling,” “April Mediated Signaling,” and “B Cell Activating Factor Signaling”). These findings are particularly striking as the physiologic levels of mechanical loading applied here (10% compressive strain) are typically associated with promoting chondrocyte anabolism (*24*), although our data is consistent with other reports suggesting a proinflammatory role of TRPV4 (*22, 54, 55*). Together, the rapidly-resolving mechanical load-induced inflammation in this context may be part of a natural cascade by which a quickly-decaying inflammatory response is characteristic of the regenerative, anabolic response to mechanical loading (*56, 57*). Additionally, increasing evidence suggests a potential role for TRPV4 in mediating cellular and tissue inflammation, and studies of chondrocyte-specific TRPV4-knockout mice report decreased severity of age-associated OA (*54*). A load-induced mechanism of chondrocyte inflammation through TRPV4 may provide a target for OA and other age-related diseases (*22, 55, 58*). The synthetic gene circuits developed in this study highlight the opportunities to target different responses through downstream pathway selection, namely using an NFKBr-circuit that is sensitive to IL-1α inflammation and a PTGS2r-circuit that is IL-1α-insensitive. To this end, the pathologic conditions present in osteoarthritis generate a milieu of inflammatory agents and factors that may, in addition to TRPV4-activation, induce NF-*κ*B signaling and *PTGS2* regulation. Deep RNA-sequencing and promotor engineering may provide a unique strategy for developing distinct, mechano-responsive tools in an inflammatory, osteoarthritic joint. Additional therapeutic targets can also be readily inhibited or activated as well; our lab has investigated using intracellular inhibitors of NF-*κ*B signaling and the soluble tumor necrosis factor receptor-1 as two alternative options for inhibiting inflammation (*59, 60*).

It is important to note some potential limitations, as well as future directions of this work. The microarray analysis offers us a first-ever look at the unique transcriptomic landscape induced by TRPV4 activation. To this end, while we used a positive strategy for the groups, i.e. comparing mechanical loading to GSK101 TRPV4 activation, using a negative strategy, i.e. comparing the mechanical loading response to mechanical loading in the presence of GSK205, may offer alternative transcriptomic targets that are TRPV4-regulated. Similarly, with the advent of next-generation sequencing technologies, performing a transcriptomic analysis of the targets of TRPV4 activation would likely increase target resolution, allowing more precise analysis of gene regulatory networks (GRNs) induced by TRPV4 activation. Subsequent GRN analysis would permit identification and discrimination between uniquely mechanically-responsive and uniquely inflammation-responsive networks to enable our long-term goals towards identifying precise genetic regulatory events driven TRPV4 activation. Next-generation sequencing technologies may also reveal more robust negatively-regulated transcript targets of mechanical loading. Our data here only uncovered several TRPV4-repressed targets, including collagen type VIII α1, suggesting more refined transcript resolution may be necessary to detect additional targets.

Using an engineered, living tissue construct for coordinated drug delivery obviates many of the traditional limitations of “smart” materials, such as long-term integration, rapid dynamic responses, and extended drug delivery without the need for replacement or reimplantation of the drug delivery system. The modular approach of using cells as both the mechanosensors and the effectors within engineered tissues allows for regulation and sensitivity at the mechanically-sensitive channel or receptor level, the signal network level, and the gene circuit level. While we used the primary chondrocyte’s endogenous TRPV4 to drive our synthetic system, engineering novel mechanically-sensitive proteins may beckon a new frontier for coordinating inputs or driving receptor activation from novel and precise mechanical inputs. Together, this framework for developing mechanically-responsive engineered tissues is a novel approach for establishing new autonomous therapeutics and drug delivery systems for mechanotherapeutics.

## Materials and Methods

### Tissue harvest, cell isolation, agarose gel casting, and culture

Full-thickness porcine articular chondrocytes were enzymatically isolated from the femurs of pigs (∼30 kg; 12-16 weeks old) using collagenase (Sigma) media. Filtered cells were mixed 1:1 with 4% molten type VII agarose (Sigma) and the cells-agarose mixture was injected into a gel apparatus and allowed to set at room temperature. Chondrocyte-laden disks were punched out yielding engineered cartilage at a final concentration of 2% agarose and 15-20 million cells/mL. All constructs were given 2-3 d to equilibrate and media were changed three times per week during chondrogenic culture using a base media which consisted of DMEM-High Glucose (Gibco) supplemented with 10% FBS (Atlas), 0.1 mM nonessential amino acids (Gibco), 15 mM HEPES (Gibco), 40 µg/mL proline (Sigma), 1x-penicillin/streptomycin, and fungizone (Gibco), and fresh 50 µg/mL ascorbyl-2-phosphate (Sigma) and maintained at 37°C and 5% CO_2_ [24]

### Chondrocyte mechanical and pharmacologic stimulation

#### Confocal imaging of mechanical compression

A custom mechanical compression device was used to compress agarose constructs while simultaneously performing intracellular calcium imaging on a confocal microscope (Zeiss LSM880) using fluo-4 AM and fura-red AM (ThermoFisher Scientific) based on manufacturer’s protocols. Opposing platens were controlled with stepper motors to apply 60 rounds of compressive loading (10%) after a 2% tare strain. Ratiometric calcium imaging (Calcium Ratio = Intensity_fluo-4_/Intensity_fura-red_) was analyzed within each sample with ImageJ for 2.5 m before and for 2.5 m after the mechanical loading.

#### High-throughput mechanical compression

A custom mechanical compression device was used for sinusoidal compression of 24-individual tissue constructs simultaneously using a closed loop displacement-feedback system [24]. This system allows for compression at 37º C and 5% CO_2_.

#### Osmotic stimulation

For calcium-imaging studies, osmotic loading media were prepared using HBSS media (Gibco). For mechanogenetic-tissue culture studies, osmotic loading media were prepared using ITS+ DMEM media. These media were titrated to hypoosmotic media by adding distilled water and measured with a freezing-point osmometer (Osmette 2007; Precision Systems). For osmotic stimulation of mechanogenetic samples, standard culture media (containing 10% FBS) was replaced with iso-osmotic (380 mOsm) ITS+ base media, containing 1% ITS+ premix (Corning), 0.1 mM nonessential amino acids, 15 mM HEPES, and 1x penicillin/streptomycin, 3 d before osmotic loading to acclimate tissues.

#### Micropipette aspiration

Detail methods for micropipette aspiration are described previously [61]. Briefly, glass micropipettes were drawn to a diameter of ∼10 µm and coated with Sigmacote (Sigma) to prevent cell binding to the glass micropipette. The micropipette was brought in contact with a cell, and a tare pressure of 10 Pa was applied for a period of 3 minutes. Increasing step pressures were then applied in increments of 100 Pa were then applied for 3 minutes each until the cell was fully aspirated. Laser scanning microscopy was used to measure cell deformation (DIC channel) and [Ca^2+^]_i_ throughout the experiment. Ratiometric calcium imaging, as described above, was performed to assess the mechanoresponse of chondrocytes to micropipette aspiration, however we found the application of step increases in pressure to the cell surface using a micropipette rarely initiated a Ca^2+^ transient.

#### Atomic Force Microscopy

Freshly isolated chondrocytes were plated on glass coverslips and incubated for 2-3 d. Prior to atomic force microscopy (AFM) compression, cells were incubated with 10 µM GSK205 or DMSO (vehicle) and stained with intracellular calcium dye Fura-2 AM (Molecular Probes) according to manufacturer’s protocols. Cells were loaded to 400 nN using a AFM (Asylum Research MFP-3D) with tipless cantilevers (k=6.2 N/m; MikroMasch) while simultaneously recording intracellular calcium levels with an Olympus microscope and cycled with 340 nm and 380 nm light to produce a ratiometric output (Calcium Ratio = Intensity_Fura-2 @ 380 nm_ / Intensity_Fura-2 $ 340 nm_) for intracellular calcium levels which were analyzed with ImageJ [44].

#### Pharmacologic TRPV4 modulation

GSK1016790A (GSK101, Millipore-Sigma) was resuspend at final concentrations (1-10 nM) and matched with a DMSO (vehicle) control. GSK205 (manufactured at the Duke Small Molecule Synthesis facility) was used as a TRPV4-specific inhibitor and pre-incubated with samples before analysis to allow for diffusion within three-dimensional tissues and used at a final concentration of 10 µM with appropriate DMSO (vehicle) controls.

#### FLIPR assay

After digestion and isolation, filtered chondrocytes were plated in a 96-well plate at 10,000 cells per well and left for 24 hours before stimulation on a fluorescent imaging plate reader (FLIPR) by which individual wells were stimulated with either osmotic or GSK101-containing media. Cellular response in each well was measured via fluo-4 intracellular calcium dye using the fluo-4 NW calcium assay kit (Molecular Probes) according to manufacturer’s directions. GSK205 was pre-incubated and added alongside stimulated and unstimulated chondrocytes to directly assess the TRPV4-dependent response of osmotic and pharmacologic stimulation.

### Finite element modeling

Finite element models of cellular deformation were performed using the FEBio (www.febio.org) finite element software package (version 2.6.4). Models were run using a Neo-Hookean elastic material for the cell and membrane compartments of the cell [62]. All model geometries utilize axisymmetry boundary conditions to reduce the model size. Osmotic loading was assessed using a Donnan osmotic loading material with parameters taken from the van’t Hoff relation for chondrocytes under osmotic loading and loading chondrocytes with a −60 mOsm osmotic media shift. Models for micropipette aspiration were run assuming a cell modulus of 1 kPa and Poissons ratio of 0.4 while imposing a pressure of −200 Pa to the cell [61]. Models of single-cell direct deformational loading (AFM) were performed by simulating an elastic sphere being compressed to 13% of its original height. FEBio testing suites were used to validate the shell, contact, and neo-Hookean code features and the Donnan model was validated against the van’t Hoff equation.

### Microarray collection and analysis

After a 14 d preculture, engineered cartilage constructs (*Ø*4 mm *×* 2.25 mm) were stimulated under 10% compressive loading or 1 nM GSK101 for 3 h/d for 3 d. An unstimulated control (free-swelling) was cultured under identical conditions. Immediately after stimulation, constructs were washed and then fed with culture media. Constructs were snap frozen in liquid N2 at 0, 3, 6, 12, 20, 24, and 72 hours after initial stimulation. Total RNA was extracted from the constructs using a pestle homogenizer and the Norgen Biotek RNA/protein purification plus kit. RNA quantity and quality were assessed by Nanodrop (Thermoscientific) and the Agilent Bioanalyzer. Total RNA was processed using the Ambion WT expression labeling kit and the Porcine Gene 1.0 ST Array (Affymetrix). The raw-signal of arrays were induced into R environment and quantile-normalized by using *‘affy’* and *‘oligo’* package. The significantly differentially expressed genes (DEGs) were identified and analyzed by using one regression model in R with package *‘genefilter’, ‘limma’, ‘RUV’, ‘splines’, ‘gplots’* and *‘plotly’*at adjusted p-value cutoff 0.05. Then the DEGs were imported into Ingenuity Pathway Analysis (IPA) to perform the pathway enrichment analysis. The DEG heatmap was plotted by using *‘gplots’* in R.

### Mechanogenetic circuit design, development, viral development, and culture

We developed two lentiviral systems consisting of an NF-κB inducible promoter upstream of either IL-1Ra (NFKBr-IL1Ra) or luciferase (NFKBr-Luc). Therefore, upon NF-κB signaling, either IL-1Ra (NFKBr-IL1Ra) or luciferase (NFKBr-Luc) is expressed as a measure of mechanogenetic circuit activation. Additionally, we developed two lentiviral systems consisting of a synthetic human *PTGS2* promoter upstream of either IL-1Ra (PTGS2r-IL1Ra) or luciferase (PTGS2r-Luc). Therefore, when *PTGS2* is activated either IL-1Ra or luciferase is expressed.

#### NFKBr-circuit design

A synthetic NF-κB inducible promoter was designed to incorporate multiple NF-κB response elements as previously described [46]. A synthetic promoter was developed containing five consensus sequences approximating the NF-κB canonical recognition motif based on genes upregulated through inflammatory challenge: *InfB1, Il6, Mcp1, Adamts5*, and *Cxcl10* [62]. A TATA box derived from the minimal CMV promoter was cloned between the synthetic promoter and downstream target genes, either murine *Il1rn* or firefly luciferase from the pGL3 basic plasmid (Promega), and an NF-κB-negative regulatory element (NRE-5’-AATTCCTCTGA-3’) was cloned upstream of the promoter to reduce background signal [46,64].

#### PTGS2r-circuit design

A synthetic human *PTGS2* promoter was obtained from SwitchGear Genomics and cloned into the NFKBr-IL1Ra or NFKBr-Luc lentiviral transfer vectors in place of the NF-κB inducible promoter. The NF-κB inducible promoter was excised using EcoRI and PspXI restriction enzymes and the *PTGS2* promoter was inserted in its place using the Gibson Assembly method to create the PTSG2r-IL1Ra and PTGS2r-Luc circuits [65].

#### Lentivirus production and chondrocyte transduction

HEK293T were co-transfected with second-generation packaging plasmid psPAX2 (No. 12260; Addgene), an envelope plasmid pMD2.G (No. 12259; Addgene), and the expression transfer vector by calcium phosphate precipitation to make vesicular stomatitis virus glycoprotein pseudotyped lentivirus (LV) [66]. The lentivirus was harvested at 24 and 48 hr post-transfection and stored at −80°C until use. The expression transfer vectors include the NFKBr-IL1Ra, NFKBr-Luc, PTGS2r-IL1Ra, and PTGS2r-Luc plasmids. The functional titer of the virus was determined with quantitative real-time polymerase chain reaction to determine the number of lentiviral DNA copies integrated into the genome of transduced HeLa cells [66]. For chondrocyte transductions, freshly isolated chondrocytes were plated in monolayer at a density of 62,000 cells/cm^2^ and incubated overnight in standard 10% FBS media. The following day, virus was thawed on ice and diluted in 10% FBS media to obtain the desired number of viral particles to achieve a multiplicity of infection (MOI) of 3 [46]. Polybrene was added to a concentration of 4 µg/mL to aid in transduction. The conditioned media of the chondrocytes was aspirated and replaced with the virus-containing media and cells incubated for an additional 24 hr before aspirating the viral media and replacing with standard 10% FBS media. Five days later, cells were trypsinized, counted, and cast in agarose as described above to prepare mechanogenetic constructs at 15 million cells/mL in 2% agarose gel. Constructs were cultured as described above until testing. Viral titers measured by qPCR revealed ∼95% of chondrocytes were transduced with this method.

#### Mechanogenetic circuit testing outcome measures

Assessing mechanogenetic tissue construct activation in IL-1Ra producing constructs was measured with an enzyme linked immunosorbent assay (ELISA) for mouse IL-1Ra (R&D Biosystems) according to manufacturer’s protocols. Data are reported as the amount of IL-1Ra produced per construct (in ng) normalized by the tissue wet weight mass of the construct (in g). Luciferase-based mechanogenetic protection was assessed using a bioluminescent imaging reader (Cytation5, BioTek) at 37 °C and 5% CO2 and cultured in phenol-red free high-glucose DMEM supplemented with 1% ITS+, 2 mM Glutamax, 1 mM sodium pyruvate, 15 mM HEPES, 40 µg/mL proline, 1x-penicillin/streptomycin, and 1 µM luciferin. Prior to stimulation, samples were imaged under free-swelling conditions for ∼1 d to get a baseline bioluminescent level (Fo). For mechanical loading studies, constructs were then mechanically loaded or similarly transferred for a free-swelling control and then returned to the bioluminescent imaging. For GSK101 pharmacologic studies, the bioluminescent media (above) was supplemented with GSK101 or DMSO (vehicle control) to the appropriate dose (1-10 nM) and simultaneously imaged for 3 h, before washing and replacing with standard bioluminescent media. Pre-stimulation baseline was normalized from post-stimulation bioluminescent readings (F) to yield F/Fo as the outcome measure.

### Coculture studies

To assess mechanogenetic anti-inflammatory protection, NFKBr-IL1Ra constructs were cultured with a porcine cartilage explant for 72 h in the presence of 0 or 0.1 ng/mL porcine IL-1α. Porcine cartilage explants (3 mm diameter) were cored from condyle cartilage and the subchondral bone was removed, leaving a cartilage explant ∼1-2 mm thick including the superficial, middle, and deep zones and cultured in iso-osmotic ITS+ media (380 mOsm, formulation listed above) base media until experimentation. Mechanical loading was applied daily with 3 h/d hypo-osmotic loading (180 mOsm) before returning to the iso-osmotic media (380 mOsm) containing the explant and IL-1α. Iso-osmotic controls were moved similarly to an iso-osmotic media (380 mOsm). Control, non-anti-inflammatory mechanogenetic tissue constructs, consisted of NFKBr-Luc tissues. Tissue sulfated glycosaminoglycan proteoglycans (S-GAG) content in the engineered tissue construct and media were assessed with the DMB assay [67]. Tissue samples were digested with papain overnight (60 °C) to measure tissue S-GAG using the DMB assay. Bulk S-GAG amount was normalized to tissue wet weight. Constructs were fixed in neutral buffered formalin overnight before embedding in paraffin and sectioning to 7 µm. Histological slices were stained with safranin-O to examine S-GAG distribution and abundance.

### Statistical analysis

Data were analyzed with one-way ANOVA or two-way ANOVA (α=0.05) where appropriate using R software (www.r-project.org). For two-way ANOVAs, individual groups were compared using a Tukey post-hoc analysis when the interaction of factors was also significant. Correlation trends of strain magnitude to IL-1Ra production was analyzed in R using linear model analysis. Temporal bioluminescent data were analyzed using a two-tailed t-test comparison between the groups at each analyzed time point.

## Acknowledgements

This work was supported by the Shriners Hospitals for Children, NIH (AR76665, AG46927, AG15768, AR74240, AR73752, AR074992, AG28716), Nancy Taylor Foundation, Arthritis Foundation, NSF EAGER Award, NSF Graduate Research Fellowship Program (DGE-1745038), and the Phillip and Sima Needleman Fellowship, the Duke School of Medicine, and a Duke Clinical and Translational Science Award (CTSA) UL1TR001117. We also thank Dr. Jonathan Brunger for his contribution in the design of the NFKBr-circuits, Holly Dressman of the Duke Microarray facility, Isaura Rigo, and Dawn Chasse for technical assistance, and Geoffrey Erickson, Scott Pritchard, and Remco Minkhorst for contributions to the cell mechanics assays.

## Author Contributions

R.J.N, L.P., and F.G. conceived the project. R.J.N, L.P., N.B.H., A.S., A.K.R., C.J.O., B.Z., A.L.M., and F.G. designed the experiments. R.J.N and A.S. conducted the chondrocyte mechanical modulation experiments and the finite element modeling. N.B.O., C.J.O., W.B.L., B.Z, and A.L.M. planned, collected, and analyzed the microarray data of TRPV4 stimulation. R.J.N., L.P., J.L., S.S., J.K., and A.K.R. conducted mechanogenetic circuit experiments and analysis. R.J.N., L.P., and F.G. wrote the manuscript. All authors read, edited, and approved the final manuscript.

## Data Availability

The microarray data generated and analyzed are publicly available (GEO in process). All additional data needed to evaluate the conclusions in the paper are present in the paper and/or the Supplementary Materials. Any other data are available from the corresponding author upon reasonable request.

## Supplementary Tables and Figures

**Supplementary Table 1**. Tabulated values of all differentially expressed genes over the first 24 h cycle after initial GSK101 stimulation taken at 0, 3, 12, and 20 h after the cessation of stimulation.

**Supplementary Table 2**. Tabulated values of all differentially expressed genes over the first 24 h cycle after initial mechanical loading stimulation taken at 0, 3, 12, and 20 h after the cessation of stimulation.

**Fig. S1.**
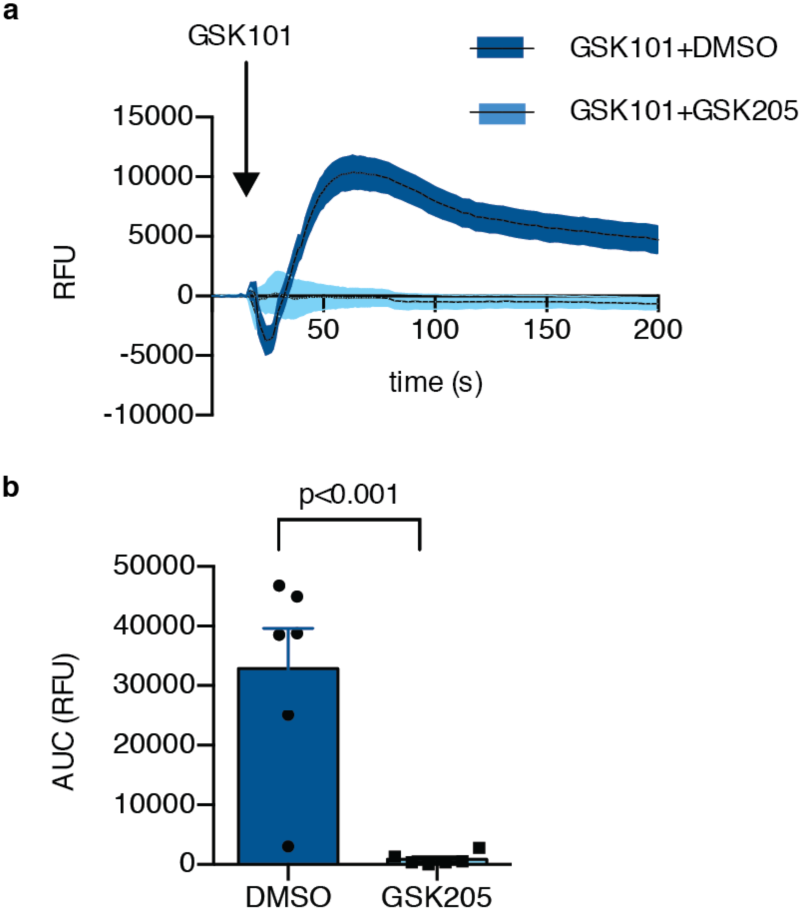
The TRPV4 agonist GSK101 stimulated intracellular calcium signaling immediately after addition (top) and this signaling was potently abolished by the antagonist GSK205. (a) Calcium signaling temporal dynamics of GSK101 activation. (b) Cumulative area-under-curve (AUC) of the temporal data (p<0.001, n=6).

**Fig. S2.**
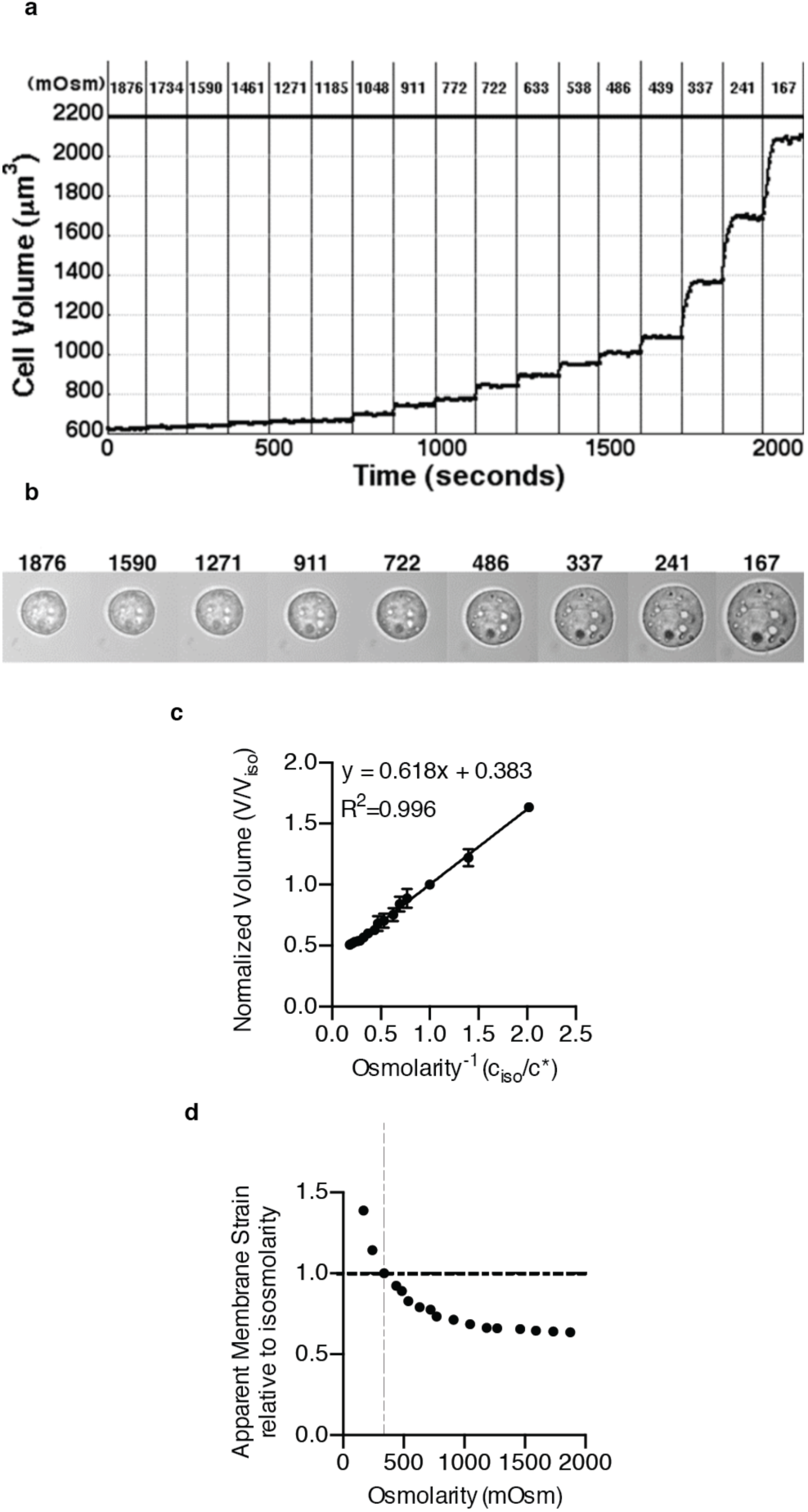
(a) Chondrocytes respond mechanically to changes in external osmolarity through a wide range of osmolarities as measured by (b) DIC images from isolated chondrocytes undergoing osmotic loading. (c) Chondrocytes behave as ideal osmometers to changes osmolarity as displayed on a Ponders plot which plots the normalized inverse osmolarity to the normalized cell volume. The slope of the line (0.618) corresponds to the volume fraction that is osmotically-active within the chondrocyte (n=18 cells). (d) Figure in (c) converted to demonstrate non-linear influence on apparent membrane strain resulting from osmotic fluctuations.

**Fig. S3.**
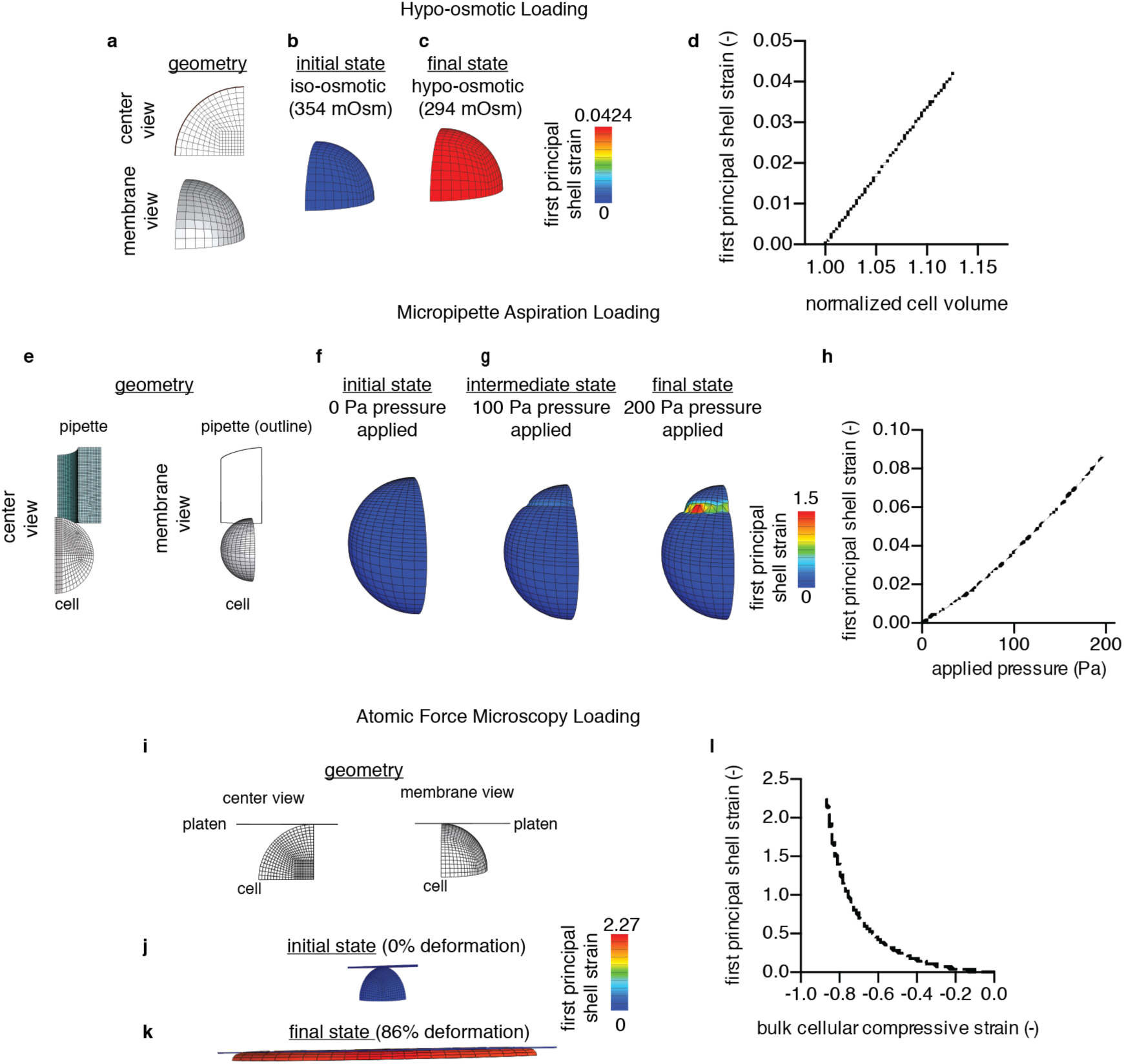
Finite element models of cellular loading conditions: hypo-osmotic (a-d), micropipette aspiration (e-h), and direct AFM loading (i-l). Models are run of an octant or quadrant cell by employing appropriate axisymmetry boundary conditions. Simulation figures represent the original geometry (a, e, i, center and membrane views), the pre-strained membrane state (b, f, j), and the deformed states (c, g, k). For osmotic loading, a hypo-osmotic step of −60 mOsm was applied; for micropipette aspiration 100 and 200 Pa pressures were applied; for AFM compression a 86% cell deformation was applied. Scale bar depicts the first principle strain of the cell membrane. Plots (d, h, l) represent the relationship between the applied deformation method and the first principal membrane strain; the micropipette plotted relationship is shown for a membrane element residing within the pipette and the AFM plotted relationship is shown for a membrane element residing along the equatorial (peak) region of membrane strain.

**Fig. S4.**
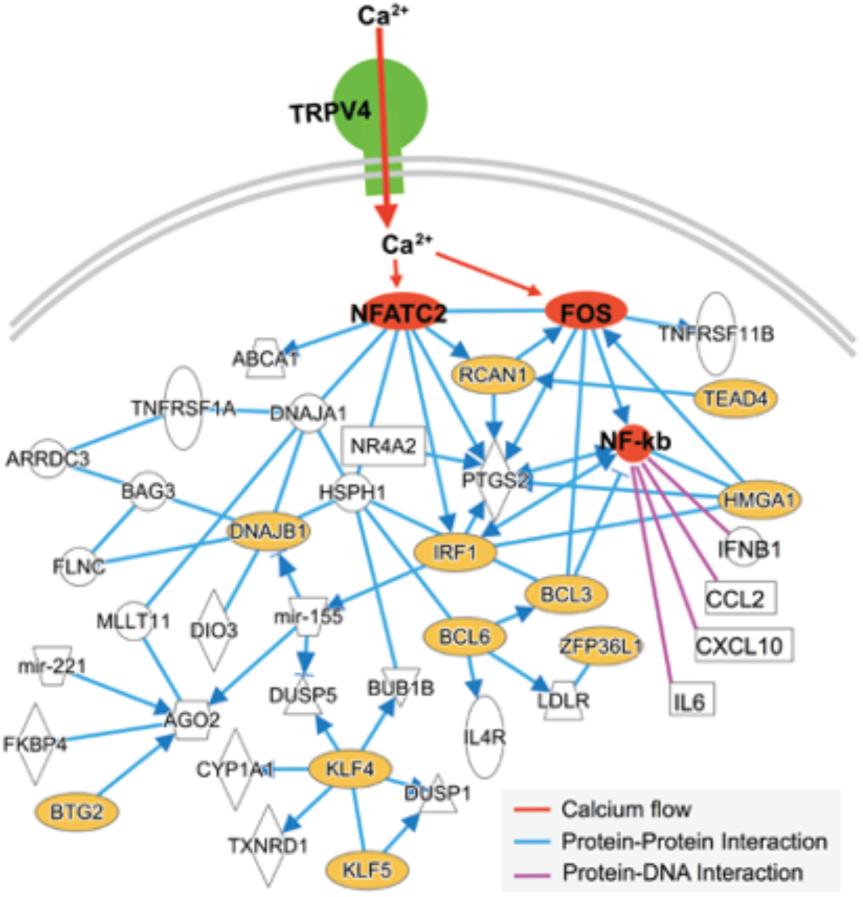
Network schematic for intracellular pathways stimulated by TRPV4 activation.

**Fig. S5.**
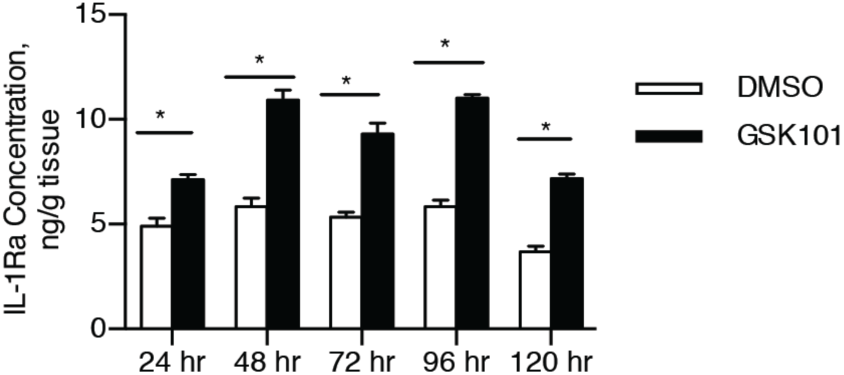
Daily IL-1Ra production from NKFBr-IL1Ra mechanogenetic cartilage exposed to either 1 nM GSK101 or vehicle (DMSO) constructs for 3 h/d every 21 h. Data are pooled and presented cumulatively in Fig. 4c. n=4 per group/timepoint, * denotes p<0.05 between GSK101 and vehicle compared on each day.

**Fig. S6.**
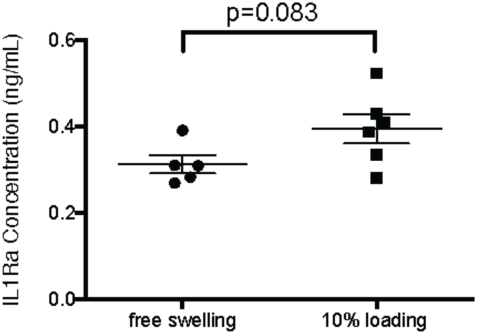
Preliminary study of PTGS2r-IL1Ra response to mechanical loading after 24 h demonstrated no significant changes in IL-1Ra levels prior to 24 h.

**Fig. S7.**
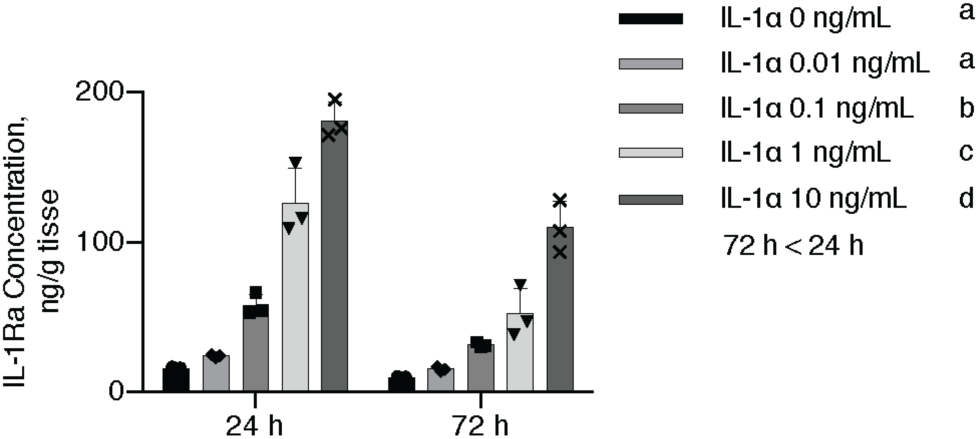
NFKBr-IL1Ra engineered cartilage responds in a dose-dependent manner to IL-1α robustly for extended durations (24 and 72 h of exposure). Subset data from 24 h exposure depicted in Fig. 5A. Different letters depict significant differences in groups.

**Fig. S8.**
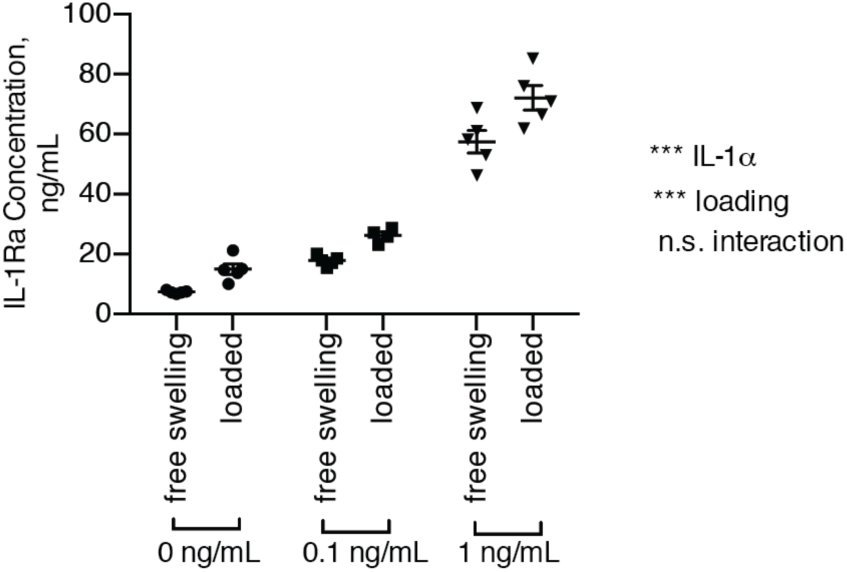
NFKBr-IL1Ra mechanogenetic tissues respond to both IL-1α and deformational mechanical loading at 24 h in a coculture system with graded IL-1α conditions (interaction n.s., p>0.05; *** denotes p<0.05).

**Fig. S9.**
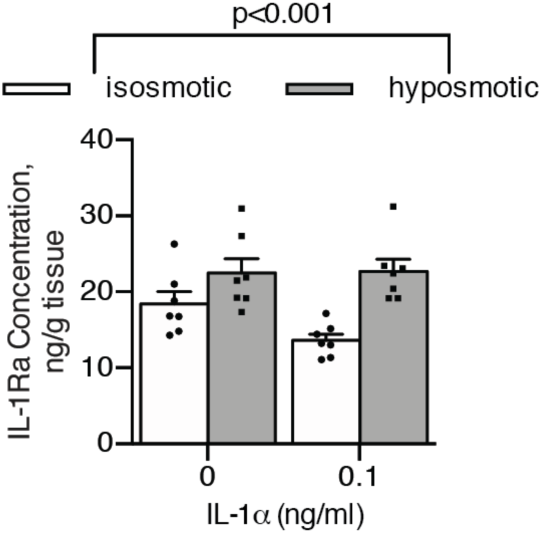
Iso-osmotic (control) and hypo-osmotic (loaded) treatments of PTGS2r-IL1Ra constructs in the presence of 0 or 0.1 ng/mL IL-1α in a coculture setting responded only to the loading factor produced a significant difference in IL-1Ra production response (p<0.001). Data are presented as mean ± SEM.

